# IMPatienT: an Integrated web application to digitize, process and explore Multimodal PATIENt daTa

**DOI:** 10.1101/2022.04.08.487635

**Authors:** Corentin Meyer, Norma Beatriz Romero, Teresinha Evangelista, Brunot Cadot, Jocelyn Laporte, Anne Jeannin-Girardon, Pierre Collet, Kirsley Chennen, Olivier Poch

## Abstract

Medical acts, such as imaging, lead to the production of several medical text report that describes the relevant findings. This induces multimodality in patient data by linking image data to free-text and consequently, multimodal data have become central to drive research and improve diagnosis. However, the exploitation of patient data is challenging as the ecosystem of analysis tools is fragmented depending on the type of data (images, text, genetics), the task (processing, exploration) and domains of interest (clinical phenotype, histology). To address the challenges, we present IMPatienT (**I**ntegrated digital **M**ultimodal **PATIEN**t da**T**a), a simple, flexible and open-source web application to digitize, process and explore multimodal patient data. IMPatienT has a modular architecture to: (i) create a standard vocabulary for a domain, (ii) digitize and process free-text data, (iii) annotate images and perform image segmentation, and (iv) generate a visualization dashboard and perform diagnosis suggestions. We showcased IMPatienT on a corpus of 40 simulated muscle biopsy reports of congenital myopathy patients. As IMPatienT relies on a user-designed vocabulary, it can be adapted to any domain of research and can be used as a patient registry for exploratory data analysis (EDA). A demo instance of the application is available at https://impatient.lbgi.fr/.

## INTRODUCTION

Patient data now incorporates the results of numerous modalities, including imaging, next-generation sequencing and more recently wearable devices. Most of the time, medical acts produce imaging data, such as echography, radiology or histology result in the production of medical reports that describe the relevant findings. Thus, multimodality is induced in patient data, as imaging data is inherently linked to free-text reports. The link between image and report data is crucial as raw images can be re-interpreted during the patient’s medical journey with new domain knowledge or by different experts leading to different reports. Thus, patient multimodal data needs to be processed in an integrated way to preserve this link in a single database.

Useful tools to centralize, process and explore multimodal data are essential to drive research and improve diagnosis. The use of multimodal data has been shown to increase disease understanding and diagnosis (Kerr et al., 2014; X. Liu et al., 2018; Venugopalan et al., 2021; Yan et al., 2019). For example, Venugopalan *et al*. integrated genetic data with image data and medical records (free-text data) to improve diagnosis of Alzheimer’s disease (Venugopalan et al., 2021). In Mendelian diseases, integration of multiple levels of information is key to the establishment of a diagnosis. For instance, in congenital myopathies (CM), a combination of muscle biopsy analysis (imaging information) with medical records and sequencing data is essential for differential diagnosis between CM subtypes (Böhm et al., 2013; Cassandrini et al., 2017; North et al., 2014). Centralization of multimodal data using dedicated software is essential to implement such an approach.

However, the ecosystem of tools for the exploitation of patient data is heavily fragmented, depending on the type of data (images, text, genetic sequences), the task to be performed (digitization, processing, exploration) and the domain of interest (clinical phenotype, histology…). Exploitation tools can be divided in two main categories: (i) tools to process the data and (ii) tools to explore the data.

Clinical reports (free-text) processing relies on the use of a standard vocabulary, such as the Unified Medical Language System (UMLS) (Bodenreider, 2004) or the Human Phenotype Ontology (HPO)(Köhler et al., 2021). Several tools have been developed to easily manage and extend these standard vocabularies, such as Protégé (Musen, 2015). Text mining processes have been developed based on these standard vocabularies, that can automatically detect keywords from free-text data. For example, Doc2HPO (C. Liu et al., 2019) can extract a list of HPO terms from free-text medical records. Other software packages, such as Phenotips (Girdea et al., 2013) have been developed to centralize and process general patient information, such as demographics, pedigree, common measurements, phenotypes and genetic results. SAMS (Steinhaus et al., 2022) and RD-Connect PhenoStore (Laurie et al., 2022) are other examples of web applications that aim to perform deep phenotyping of patients by building a single database of standardized patient data using well-established ontologies sur as HPO. Finally, for imaging data, software to process and annotate gigapixel-scale microscopy images are widely used, including Cytomine (Marée et al., 2016), SlideRunner (Aubreville et al., 2018) and Ilastik (Berg et al., 2019). Cytomine is a powerful software package for gigapixel scale image annotation and analysis, that includes an ontology builder and complex image processing tools. However, it is restricted to image data only.

A wide range of tools have been developed to analyze and explore patient data. For example, based on a list of HPO terms describing a patient’s specific phenotypic profile, Phenolyzer (Yang et al., 2015) and Phenomizer (Köhler et al., 2009) can be used to help prioritize candidate genes or rank the best-matching diseases. However, these tools are restricted to the use of HPO terms to describe the patient’s profile and are not compatible with other ontologies. Ontology agnostic algorithms have also been developed that predict an outcome based on a list of terms from any normalized vocabulary, such as the Bayesian Ontology Query Algorithm (BOQA) (Bauer et al., 2012). For patient images exploitation, guidelines and frameworks have been proposed to standardize the measurement of pathological features from DICOM lung images (Cinaglia, Tradigo, et al., 2018).

Some multimodal approaches such as ClinPhen (Deisseroth et al., 2019) and Exomiser (Smedley et al., 2015) have successfully combined multiple levels of information with both phenotype information (HPO terms) and genetic information (variants) to rank candidate genes in Mendelian diseases. Other tools such as INTEGRO (Cinaglia, Guzzi, et al., 2018) have been developed to automatically data-mine disease-gene associations for a specific input disease from multiple curated sources of knowledge.

This large ecosystem of tools highlights the need for an integrated tool that can: (i) both process and explore patient data, (ii) manage multimodal data (text and images), and (iii) work in any domain of interest.

In this study, we present IMPatienT (**I**ntegrated digital **M**ultimodal **PATIEN**t da**T**a), a free and open-source web application that aims to be an integrated tool to digitize, process and explore multimodal patient data. IMPatienT is a turnkey solution that aims to aggregate patient data and provides simple tools and interfaces for a clinician to extract information from multimodal patient data in a single endpoint. Using a modular architecture, we developed four components to: (i) create a standard vocabulary describing a domain of interest, (ii) digitize and process free-text records by automatically mapping them to a set of standard terms, (iii) annotate and segment images with standard vocabulary, and (iv) generate a dashboard with automatic visualizations to explore the patient data and perform automatic diagnosis suggestions.

Finally, we demonstrate the usefulness of IMPatienT on a set of congenital myopathy (CM) cases. CM are a family of rare genetic diseases, including multiple distinct subtypes, that still lack proper diagnosis with more than 50% of patients without a genetic cause identified(H et al., 2018). We exploited IMPatienT to create a list of standard muscle-histology terms that were then used to process patient histological records and annotate biopsy images. Finally, multiple exploratory visualizations were automatically generated.

## MATERIALS AND METHODS

IMPatienT is a web application developed with the Flask micro-framework, which is a Python-based web framework. Figure 1 illustrates the global organization of the web application. The web application is composed of four modules: (i) Standard Vocabulary Creator, (ii) Report Digitization, (iii) Image Annotation, and (iv) Automatic Visualization Dashboard. All modules incorporate free, open-source and well-maintained libraries that are described in detail in the corresponding sections.

**Figure 1:**
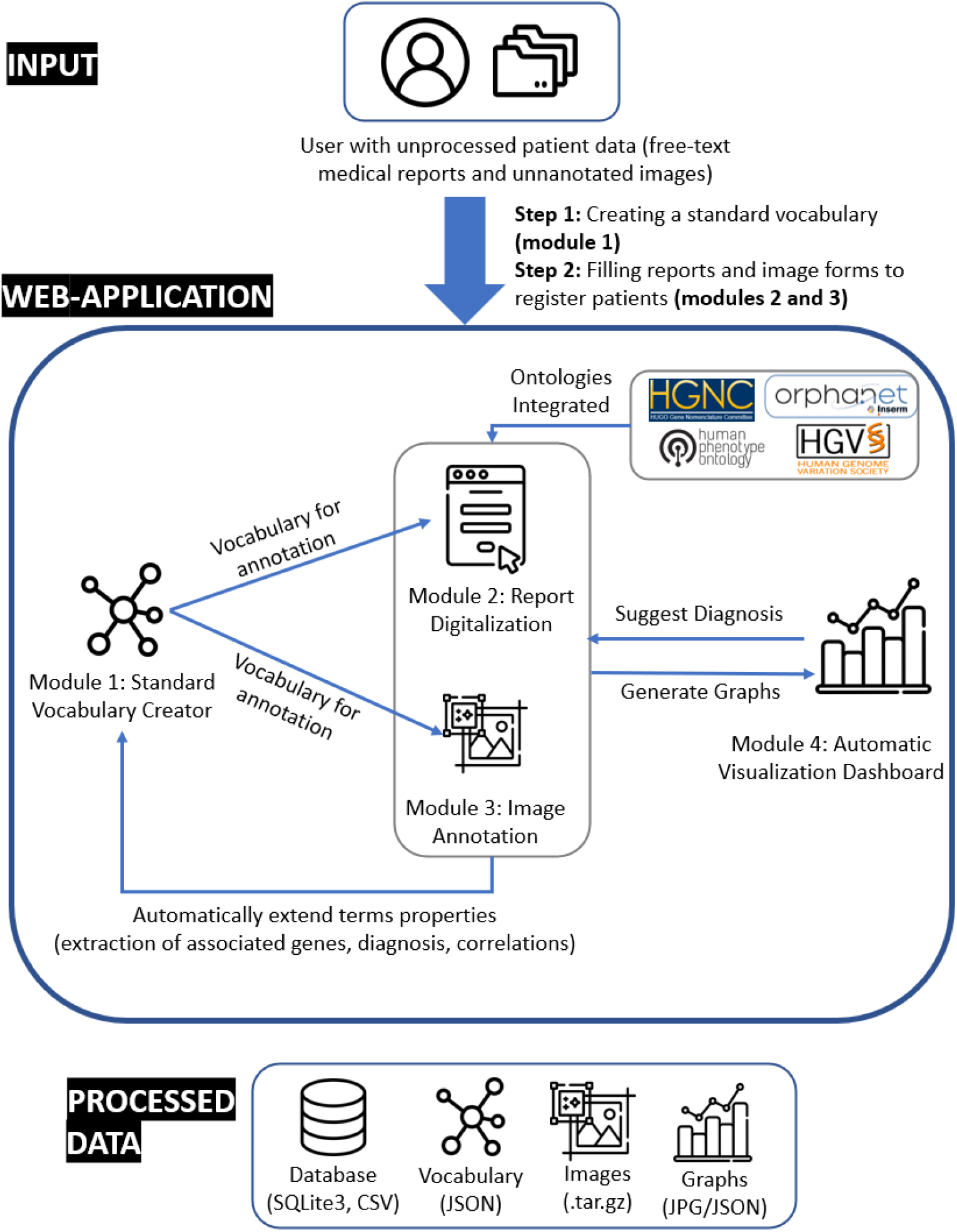
IMPatienT web application organization

### Module 1: Standard Vocabulary Creator

The standard vocabulary creator module allows to create and modify a hierarchical list of vocabulary terms with rich definitions that can be used as an image annotation class, for text reports processing, or suggestion of diagnosis. The standard vocabulary is an essential module of IMPatienT as it interacts with all subsequent modules.

Figure 2 shows a screenshot of the page used to create and manage the standard vocabulary tree. The ergonomic drag and drop system using the graphical user interface (GUI) allows the user to intuitively and quickly edit and reorganize the vocabulary to add new terms or modify existing ones. Also, the vocabulary term (node) detailed form makes it easy to edit term properties.

**Figure 2:**
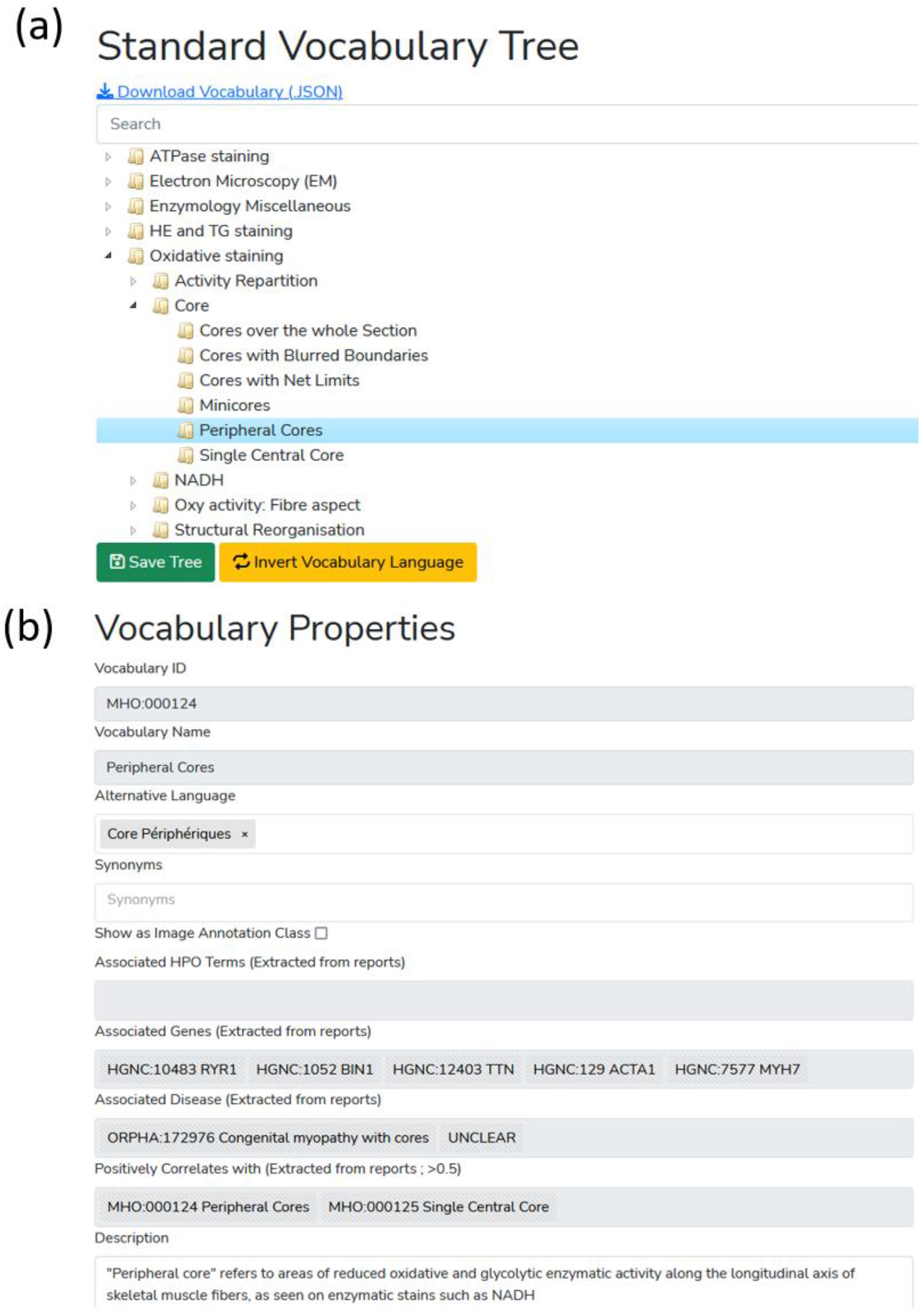
Screenshot of the Standard Vocabulary Creator module (module 1). **(a)** The hierarchical structure viewer and editor tool that supports drag and drop modification and creation/deletion/modification using the mouse. **(b)** The properties of the selected term node with its unique ID, display name, alternative language translation, synonyms, description, associated genes and diseases and correlating terms extracted from the application instance database.

The tree is generated and rendered with the JavaScript library JSTree (version 3.3.12). Each node (term) can have only one parent. For each created node (vocabulary terms), the user can assign a name and organize the tree structure (hierarchy) through the drag and drop interface. Each term in the tree is associated with nine optional properties. Four properties are defined by the user: description, list of synonyms, translation in another language, show the term as annotation class. Two properties are automatically generated: the term’s unique identifier (ID) and the hexadecimal color associated with the term (for image annotation). Additional term properties (associated diagnosis/disease class, associated genes, list of positively correlating terms [*i*.*e*. co-occurring terms in reports]) are extracted from patient records registered in the database.

Finally, if the user defines an alternative translation for terms, there is an “invert vocabulary language” button to conveniently switch between standard vocabulary languages. For instance, the user can create a vocabulary in any language and define the translation in English, then switch between the two display modes easily.

### Module 2: Report Digitization

The standard vocabulary terms are used to process documents that are in a free-text format. Module 2 uses a semi-automatic approach for digitization and processing of free-text reports that combines fast automatic detection of terms with manual reviewing of the detection. The interface of Module 2 is a form divided into four parts (Figure 3).

**Figure 3:**
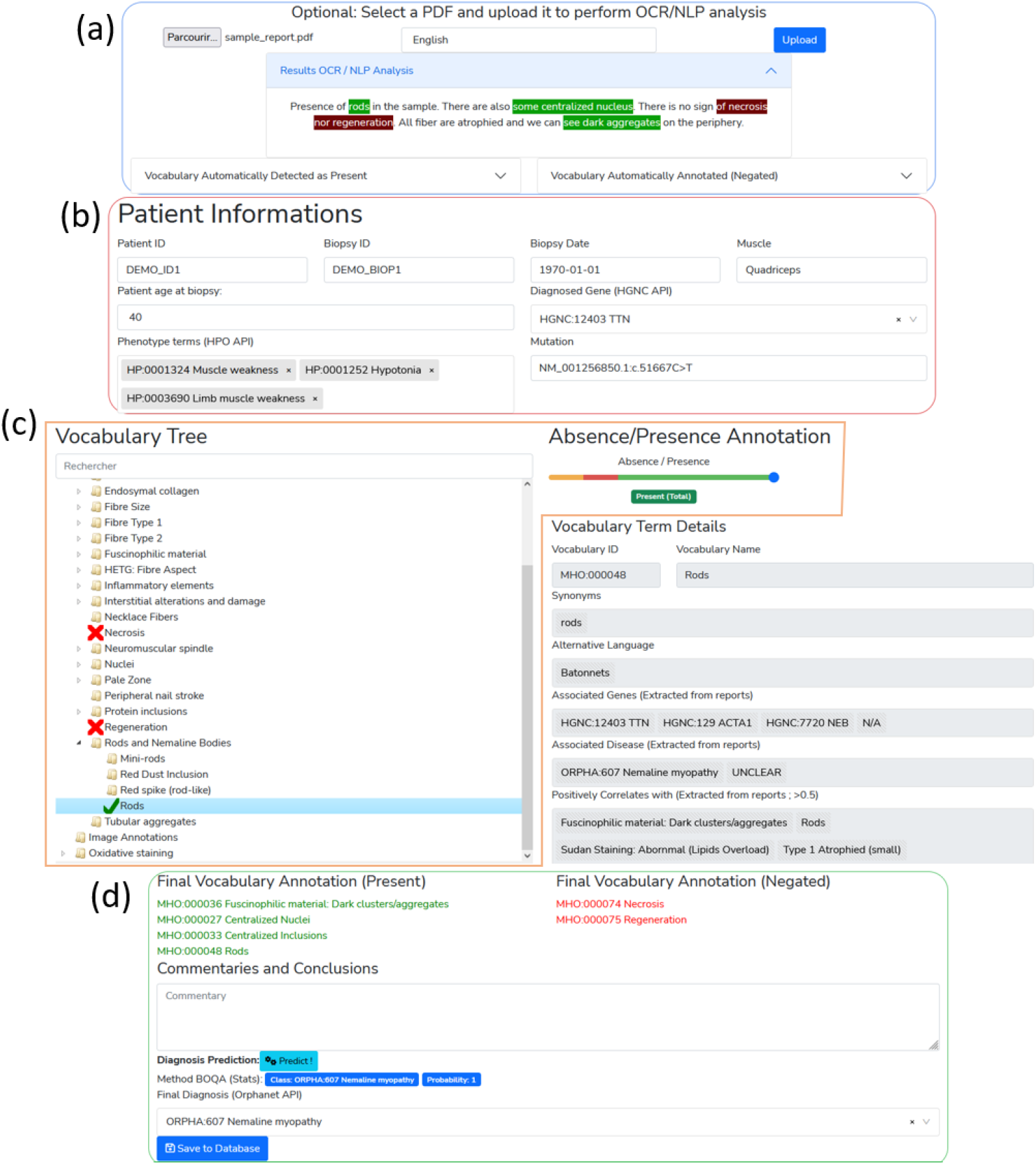
Screenshot of the report digitalization module. **(a)** PDF upload section for automatic keyword detection in the text. Detected keywords have a green background, detected and negated keywords have a red background. **(b)** Patient information section (age, document ID, gene, mutation, phenotype). **(c)** Standard vocabulary tree viewer to select keywords with associated slider to manually indicate keyword value (absence or presence level). Keywords marked as present are indicated with a green check mark, absent keywords are marked with a red cross. **(d)** Final section with an overview of all annotated terms, diagnosis selection and commentary part with automatic diagnosis suggestion using BOQA algorithm.

In the first part of the digitization form (Fig 3a), a PDF file of the free-text report can be uploaded for natural language processing (NLP) of the content. The text of the PDF report is automatically extracted and processed with NLP. The NLP method is only used to detect histological terms defined in the standard vocabulary. Detected standard vocabulary terms are highlighted (see corresponding section below “Optical Character Recognition and Vocabulary Terms Detection”). Highlighted terms allow to easily identify what standard vocabulary terms were detected as present or in negative form. This is useful for quantitative performance assessment.

The second part (Fig 3b) of the digitization form contains patient informations, such as patient ID, document ID, age of the patient. This section also allows the user to input patient informations that are not defined by the standard vocabulary and thus, not processed in the NLP section. For example, IMPatienT exploits well-established ontologies to normalize the genetic diagnosis and phenotypes (Fig 4). For example, in the gene field, when the user input characters, gene symbols are retrieved from the HUGO Gene Nomenclature Committee (HGNC) and suggested.(Tweedie et al., 2021) Mutation notations are formatted according to the Human Genome Variation Society (HGVS) sequence variant nomenclature(den Dunnen et al., 2016). Phenotypes are retrieved and suggested using the HPO ontology. None of these fields contain patient-identifying data and are optional.

**Figure 4:**
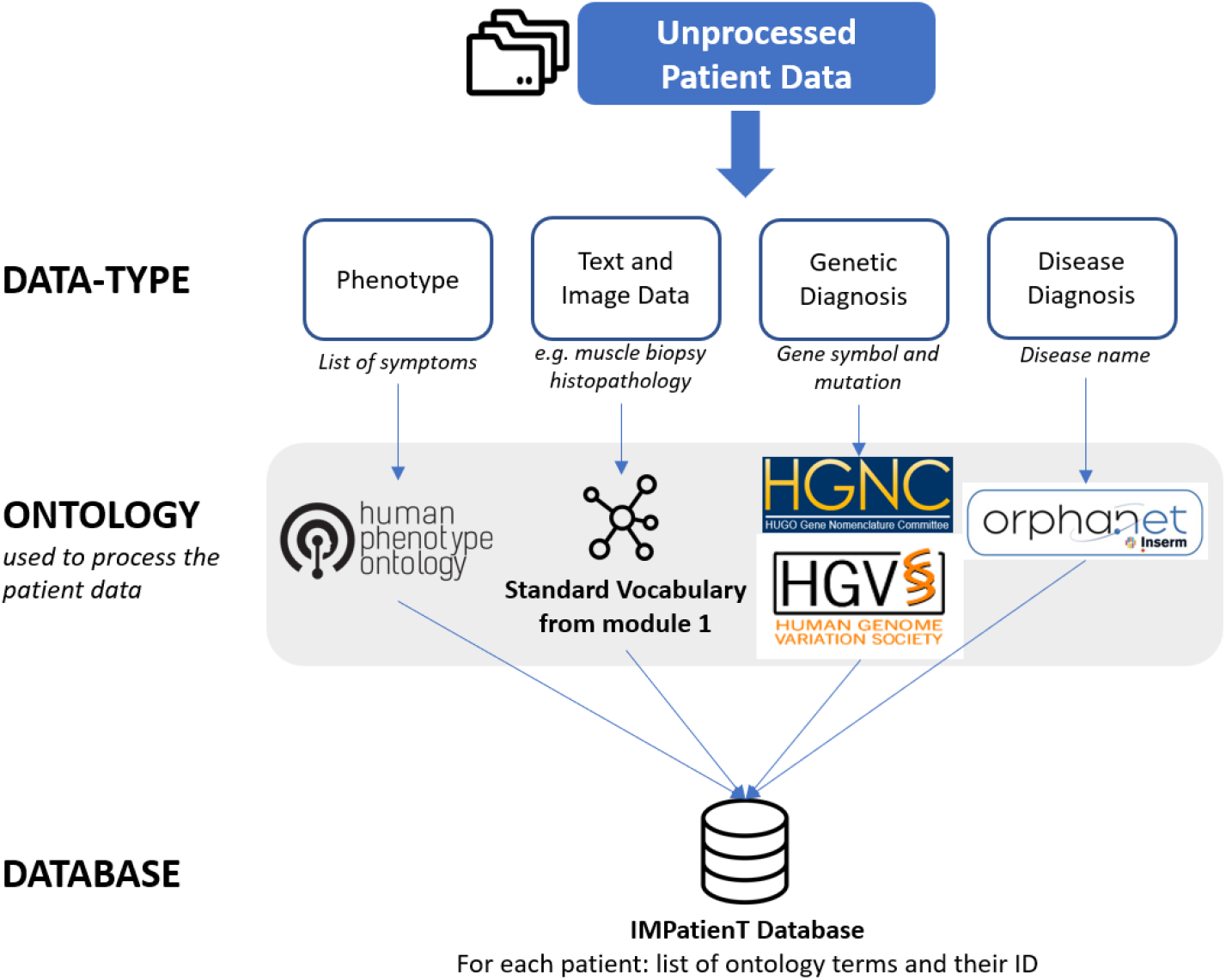
Overview of the ontologies used by IMPatienT to process patient data in the report digitization module (module 2).

The third part of the digitization form (Fig 3c) contains the standard vocabulary tree viewer with an absence/presence slider. This section allows the user to correct the automatic detection of the NLP method or to add new observations. Each vocabulary term can be marked as present, absent or no information. For terms marked as present, the slider is used to indicate a notion of quantity or certainty of the term. For example, the statement “There are a small number of fibers containing rods” can be annotated by hand by setting the vocabulary “Rods” to the value “Present” with a low quantity value. For terms that have been automatically detected, this slider value is automatically set to 0 (present in a negated sentence) or 1 (present).

Finally, the fourth part (Fig 3d) of the form allows the user to input comments and a final diagnosis for the patient, disease name are suggested from the Orphanet (INSERM, 1997) knowledge base. It also includes an automatic suggestion of the diagnosis based on already registered patients using BOQA (Bauer et al., 2012) (see the corresponding section below “Patient Disease Suggestions Method”).

### Optical Character Recognition and Vocabulary Term Detection

The patient report digitization in module 2 is facilitated by the automatic text recognition and keyword detection method. The user uploads a PDF version of the text reports to perform Optical Character Recognition (OCR), followed by Natural Language Processing (NLP) to automatically detect terms from the standard vocabulary in the report. The NLP method is only match the raw text to the standard vocabulary defined in Standard Vocabulary Module 1. Figure 5 describes the workflow of the vocabulary terms detection method. First the PDF file is converted to plain text using the Tesseract OCR (implemented in python as pyTesseract). Then, the text is processed with Spacy, an NLP python library, by splitting the text into sentences and then into individual words. The resulting list of sentences is then processed to detect negation using a simple implementation of the concept of NegEx (Chapman et al., 2001). An *n*-gram (monograms, digrams, and trigrams) procedure is applied to the list of words to identify contiguous words in the context of all the sentences of the report. The *n*-grams are then mapped against the user-created standard vocabulary using fuzzy partial matching (using Levenshtein distance) with a score threshold of 0.8. Matched keywords are kept and shown on the interface with a green or red highlight of the detected text using Mark.JS JavaScript library (green indicates the presence of the keyword, red indicates the presence in a negated sentence). Keywords are also automatically marked as present or absent (negated) in the vocabulary tree.

**Figure 5:**
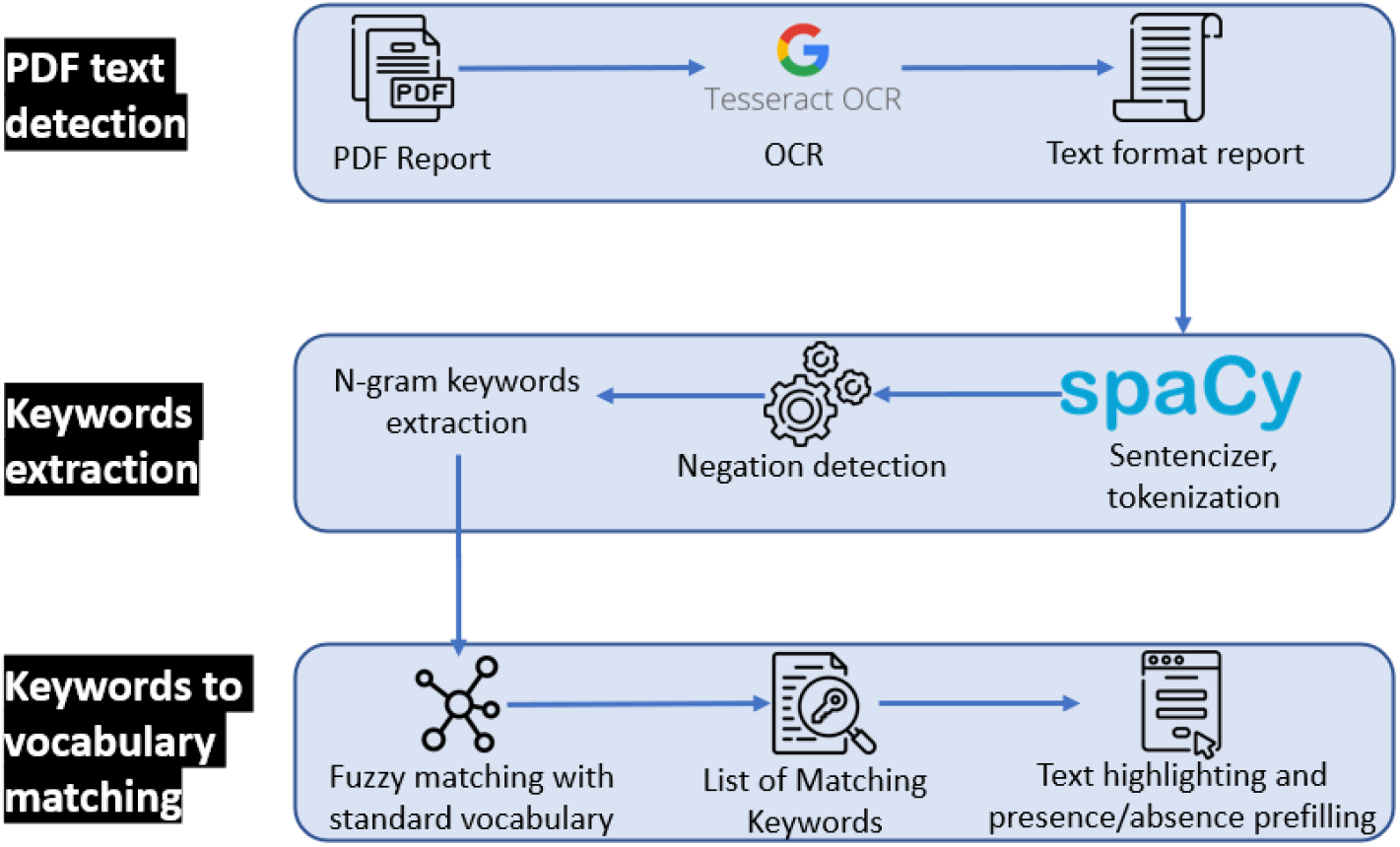
Optical character recognition and vocabulary term detection method used in the report digitization module (module 2) to automatically analyze free-text reports.

### Disease Suggestions

The report digitization module 2 contains a disease recommendation algorithm inspired by the BOQA algorithm described by Bauer *et al*. (Bauer et al., 2012). Basically, the algorithm computes the similarity between a list of input vocabulary terms annotated as “present” for a patient (the query) and a simulated patient profile for each disease class (model report) that is generated based on the data from already registered patients.

We implemented this algorithm in python, and we modified it to use the frequencies of vocabulary terms per disease for the generation of the model report instead of the initial deterministic way (not frequency aware). This means that the model report is generated based on the probability (frequency) of each vocabulary term. For example, if disease A is annotated with vocabulary term B at a frequency=0.9 and vocabulary term C at a frequency=0.1, the generated model report for disease A will have a probability=0.9 of containing vocabulary term B and a probability=0.1 of containing vocabulary term C.

Due to the stochastic nature of the generation of the model report, for any given prediction, the generation and computation of the similarity with the query is repeated 50 times. For each repetition, if a disease has a prediction probability>0.5, it is considered to be the best prediction, otherwise the prediction is “no prediction”. Finally, of the 50 repetitions, the prediction with the highest occurrence is taken as the final prediction.

### Module 3: AI-Assisted Image Annotation Using Automatic Segmentation

To process patient image data, we developed the image annotation module (module 3) to upload, annotate and perform image segmentation with standard vocabulary terms. This module is based on the *“interactive image segmentation with Dash and Scikit-image”* demonstration application (Gouillart, 2020; Hossain, 2019; Walt et al., 2014). The original source code was modified to be compatible with the standard vocabulary tree and the database.

The interactive interface to annotate image features with standard vocabulary terms is presented in figures 6a and 6b. The interface allows the user to draw a free-shape area (annotation) associated with a standard vocabulary term (class). Then, with a minimal number of user annotations, the whole image is segmented based on the annotations (shapes) provided by the user.

**Figure 6:**
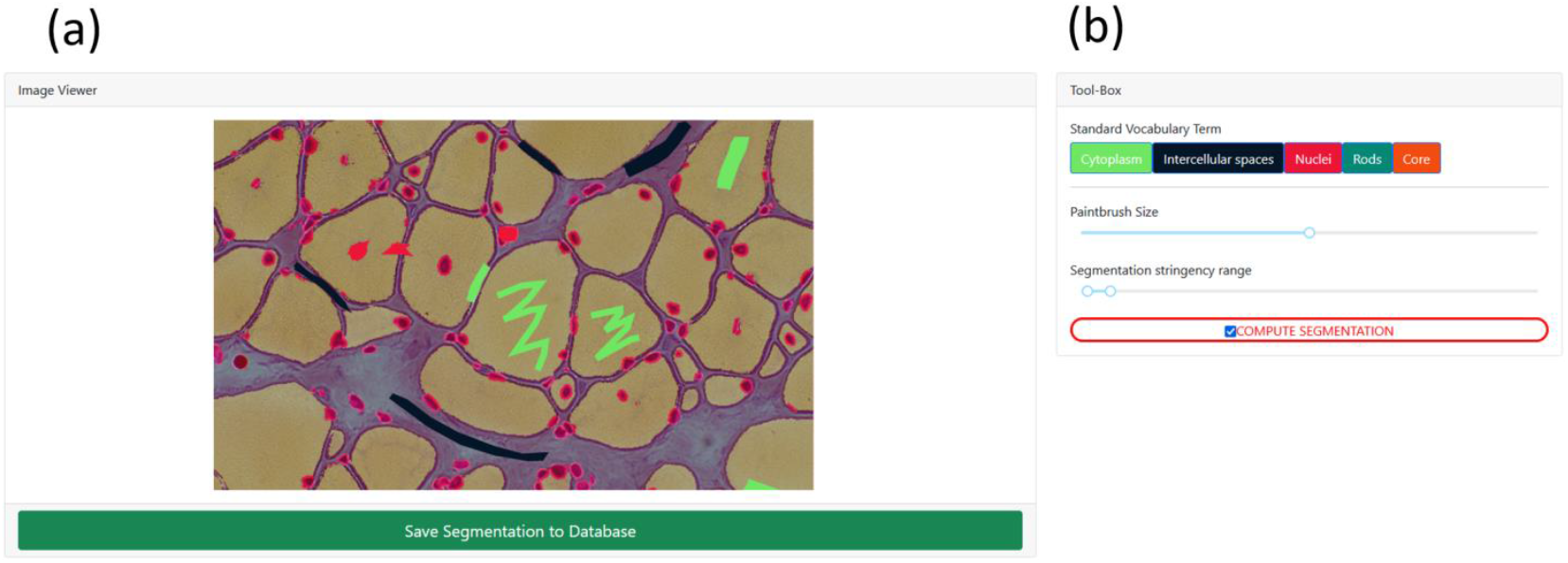
Screenshot of the image annotation module. **(a)** Image viewer used to navigate, zoom and annotate the histology image. **(b)** Menu interface to select the annotation label, brush width and segmentation parameters.

To perform image segmentation, on the server side, local features (intensity, edges, texture) are extracted from the labeled areas of the image and are used to train a dedicated AI random-forest classifier model. This dedicated model is then applied to predict similar areas in the whole image. Finally, every pixel of the image is labeled with a standard vocabulary term corresponding to the AI prediction based on the annotations. The segmentation is entirely interactive. After the initial segmentation, the user can correct the classification by adding more annotation shapes to the image and can modify the paintbrush width setting to make more precise annotation marks. In addition, the stringency range parameter of the model can be adapted using the slider to modify the model behavior and automatically recompute the segmentation in real time.

Results of the segmentation are retrievable as a single archive including the raw image, the annotations (JSON), the random-forest trained classifier, the blended image and the segmentation mask image.

### Module 4: Automatic Visualization Dashboard

The automatic visualization dashboard module is designed to perform exploratory data analysis by generating multiple graphs based on the patient data in the database. All visualizations are created using Plotly, a python graph library, that allows making interactive graphs.

### Interaction Between the Modules

IMPatienT is divided into four modules that are interconnected. The standard vocabulary module provides the vocabulary used for the image annotation module and for the NLP method used for the (histologic) standard vocabulary terms detection in the report digitization module. Any modification in the vocabulary is automatically propagated to these modules, updating the form templates and triggering the recalculation of all visualizations with the latest vocabulary information. Any modification to the standard vocabulary also updates all patients in the database to the latest version of the vocabulary, meaning that term names and definitions will be updated, and deleted terms will be marked as outdated. Adding patient information in the database, whether they are text reports (module 2) or images data (module 3), will automatically update the visualization dashboard with the latest patient information in the database. The term frequency statistics calculated by the visualization dashboard and used by the disease suggestion algorithm are automatically updated as well, providing live performances increase. The visualization dashboard is also directly linked to the standard vocabulary and during the generation of the visualizations, the rich definition of the standard terms is updated with newly associated genes, diagnosis and positively correlating terms.

### Application Security and Personal Data

IMPatienT is developed as a free and open-source project meaning that the code can be audited by anyone in the GitHub code repository. The code is regularly scanned for known issues and outdated libraries to mitigate security issues. There is no patient-identifying data kept in the database, only a custom identifier and age. No name or date of birth is required or stored.

Additionally, access to all modules and data entered via the web application is restricted by a login-page and user accounts can only be created by the administrator of the platform. No user information is stored except for the username, email and salted and hashed passwords.

## RESULTS

IMPatienT is an interactive and user-friendly web application that integrates a semi-automatic approach for text and image data digitization, processing, and exploration. Due to its modular architecture and its standard vocabulary creator, it has a wide range of potential uses.

### IMPatienT Main Functionalities

Table 1 shows the main functionalities of IMPatienT compared to other similar tools used in the community. IMPatienT integrates tools that are simple, portable, easy to implement and similar to multiple state-of-the-art solutions but in a single platform. Out of 18 selected features, IMPatienT integrates 14 of them versus a mean of 4.4 for other software with the best one being SAMS and PhenoStore integrating 6 features each. However, software such as SAMS, PhenoStore, Phenotips and Cytomine each integrates features that are not yet present in IMPatienT.

**Table 1:** Comparison of functionalities from IMPatienT compared to common state-of-the-art tools.

| Group | Functionalities | IMPatientT | Phenotips | PhenoStore | SAMS | Protégé | Doc2HPO | Cytomine | Ilastik | INTEGRO |
| --- | --- | --- | --- | --- | --- | --- | --- | --- | --- | --- |
| General Application Characteristics | Web application | X | X | X | X |  | X | X |  |  |
|  | Patient database | X | X | X | X |  |  |  |  |  |
|  | Free to use and open-source | X |  |  | X | X | X | X | X | X |
|  | Support multimodal data | X |  |  |  |  |  |  |  |  |
|  | Support for patient pedigree data |  | X | X |  |  |  |  |  |  |
| Standard Vocabulary | Vocabulary Builder | X |  |  |  | X |  | X |  |  |
|  | Advanced vocabulary terms definition | X |  |  |  | X |  |  |  |  |
|  | Full-featured ontology builder |  |  |  |  | X |  |  |  |  |
| Report digitization | Integrates reference ontologies (HPO, Orphanet) | X | X |  | X |  | X |  |  | X |
|  | Form for patient medical report digitization | X |  | X | X |  |  |  |  |  |
|  | Text recognition with OCR | X |  |  |  |  |  |  |  |  |
|  | Text processing with NLP | X |  |  |  |  | X |  |  |  |
|  | Export data to Phenopacket format |  |  | X | X |  |  |  |  |  |
| Image annotation | Image annotation and segmentation with AI | X |  |  |  |  |  | X | X |  |
|  | Support for DICOM and whole slide images |  |  |  |  |  |  | X |  |  |
| Patient data exploitation | Automatic visualization dashboard | X |  | X |  |  |  |  |  |  |
|  | Diagnosis prediction system | X | X |  |  |  |  |  |  |  |
|  | Data mining of information for specific diagnosis | X |  |  |  |  |  |  |  | X |

IMPatienT implements novel functionalities to process and exploit patient data. For example, IMPatienT is compatible with any domain of research thanks to its standard vocabulary builder. Also, with the OCR/NLP method, IMPatienT can process histologic text reports, allowing the user to exploit scanned documents. Finally, IMPatienT also provides useful utilities to exploit patient data with the various visualizations, the term, frequency table, correlation matrix and the automatic enrichment of the vocabulary terms definition (associated genes and diseases).

### IMPatienT Usage

Figure 1 shows how the user can interact with the web application to digitize, process, and explore patient data. In IMPatienT, modules can be used independently, allowing users to only use the tools they need. For example, a user might only have text report data, in this case they would be able to use the standard vocabulary creator, the report digitization tools and the visualization dashboard to process and explore their data. In another scenario, a user could only be interested in annotating an image dataset using a shared standard vocabulary that can be modified and updated collaboratively. In this use case, they would be able to only use the standard vocabulary creator and the image annotation module. However, the main strength of IMPatienT lies in the multimodal approach it provides and the module interactions.

For the complete multimodal approach, the first step is to create a standard vocabulary using the Standard Vocabulary Creator interface (module 1). The user only needs to create a few terms (nodes) to begin using the web application. Defining the properties of the terms (definition, synonyms…) is optional, and organizing them in a hierarchical structure is also optional.

Then, the user can start digitizing patient reports using module 2 (step 2). This can be done manually by filling out the form in module 2 and checking terms as present or absent in a given report, or the user can employ the Vocabulary Term Matching method by uploading a PDF version of the report. Using module 3, the user can also upload, annotate, and segment image data.

Finally, the user can view multiple exploratory graphs (histograms, correlation matrix, confusion matrix, frequency tables) that are automatically generated in module 4. All data entered via the web application are retrievable in standard formats, including the whole database of reports as a single SQLite3 file or CSV files, the images and their segmentation models and masks as a GZIP archive, the standard vocabulary with annotation as a JSON file and various graphs and tables as JSON or PNG files.

### Use Case: Congenital Myopathy Histology Reports

As a use case of IMPatienT, we focused on congenital myopathies (CM). We used the standard vocabulary creator to create a sample muscle histology standard vocabulary based on common terms used in muscle biopsy reports from the Paris Institute of Myology. Then, we inserted 40 generated digital patients in the database with random sampling of standard vocabulary terms and associated a gene and disease class among a list of common CM genes and three recurring CM subtypes (nemaline myopathy, core myopathy and centronuclear myopathy). All these data are available on the demo instance of IMPatienT (https://impatient.lbgi.fr/).

For text data, Supplementary Figure S1 shows the results of the automatic NLP method applied to an artificial muscle histology report. Twenty-two keywords were detected and match to the standard vocabulary and seven of them were detected in negated sentences (red highlight). Among the 22 vocabulary terms detected. Out of the twenty-two keywords, eighteen were correctly detected and one was detected in the wrong state of negation: “abnormal fiber differentiation” is highlighted as negated while it is present is a non-negated sentence part. Three keywords (fiber type, internalized nuclei, centralized nuclei) were detected as matching for multiple keywords from the vocabulary at the same time due to high similarity. For example, the keyword “internalized nuclei” and “centralized nuclei” have a similarity score of 86 using the Levenstein distance. Two keywords defined in the standard vocabulary were missed and not highlighted: “biopsy looks abnormal” (“abnormal biopsy” in the vocabulary) and “purplish shade” (“purplish aspect” in the vocabulary).

For the image data, figure 7 shows an example of the segmentation of a biopsy image, where we annotated the cytoplasm of the cells (green), intercellular spaces (black) and cell nuclei (red). The raw image (Fig 7a) is annotated with free-shape areas associated with standard vocabulary terms (Fig 7b). Then, the whole image is automatically segmented based on the annotations, producing the segmentation mask where each pixel is associated with a class (Fig 7c 7d).

**Figure 7:**
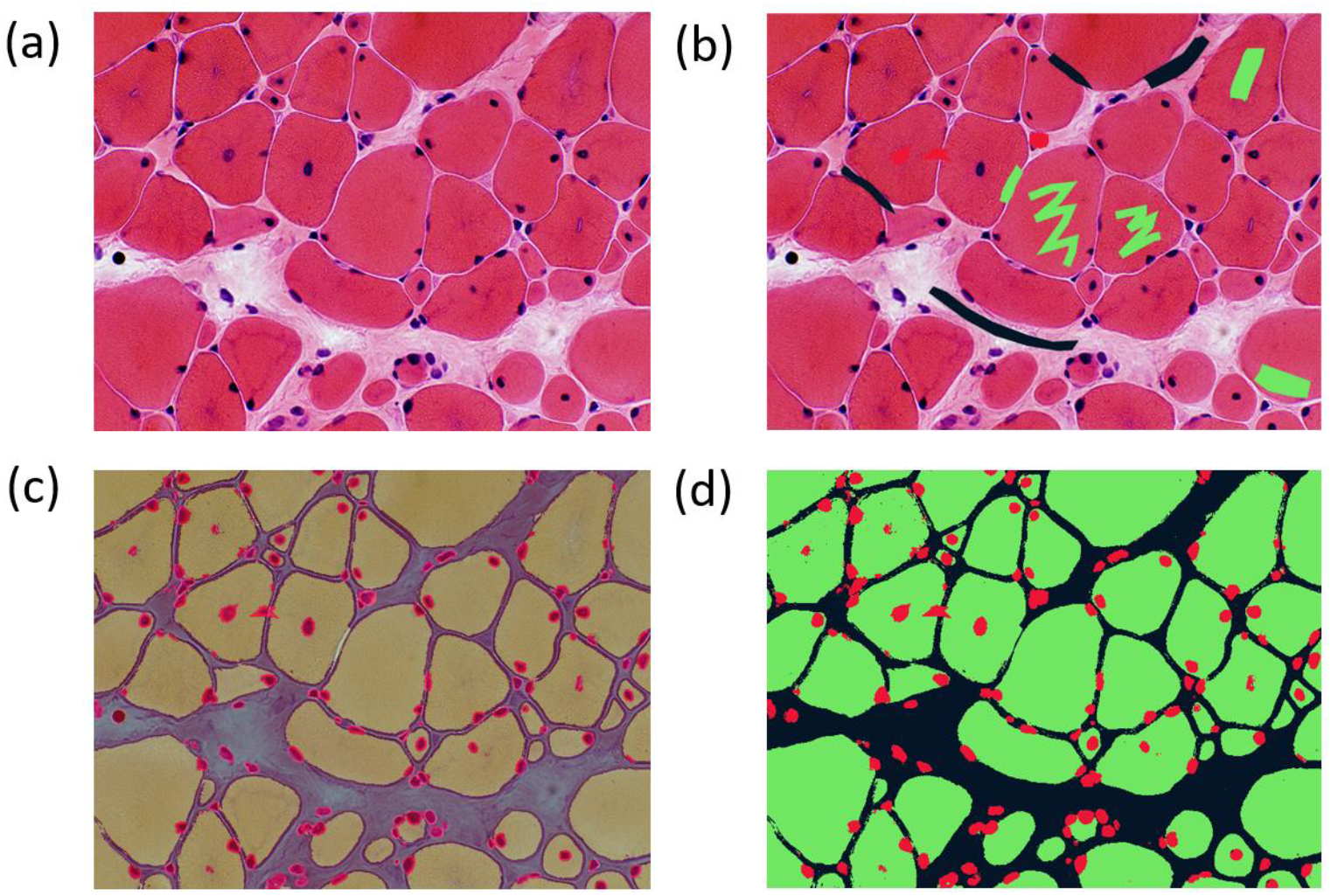
Image segmentation process in the image segmentation module. **(a)** Raw image input before annotation. **(b)** Image with limited manual annotation of cytoplasm (green), cell nucleus (red) and intercellular space (black). **(c)** Blended image of the raw image and segmented image after automated segmentation with a random-forest classifier. **(d)** Segmented image mask alone.

The automatic visualization dashboard was used to generate the six visualizations provided in figure 8. These visualizations include a breakdown of the patients in the database by age, genes, or diagnosis (Fig 8a). A correlation matrix (using Pearson correlation coefficient) between the occurrence of standard vocabulary terms is generated (Fig 8b), which can serve as a starting point for exploration of co-occurrence of features in patients. The confusion matrix of the final diagnosis of patients versus the suggested diagnosis with BOQA (Fig 8c) allows the user to monitor the accuracy of the disease suggestion function. In addition, a histogram showing the classification of patients without a final diagnosis is provided to indicate possible prognosis of undiagnosed patients (Fig 8d). Finally, the frequency of each standard vocabulary term by gene and by disease is automatically calculated and shown in two tables (see supplementary tables S1 and S2).

**Figure 8:**
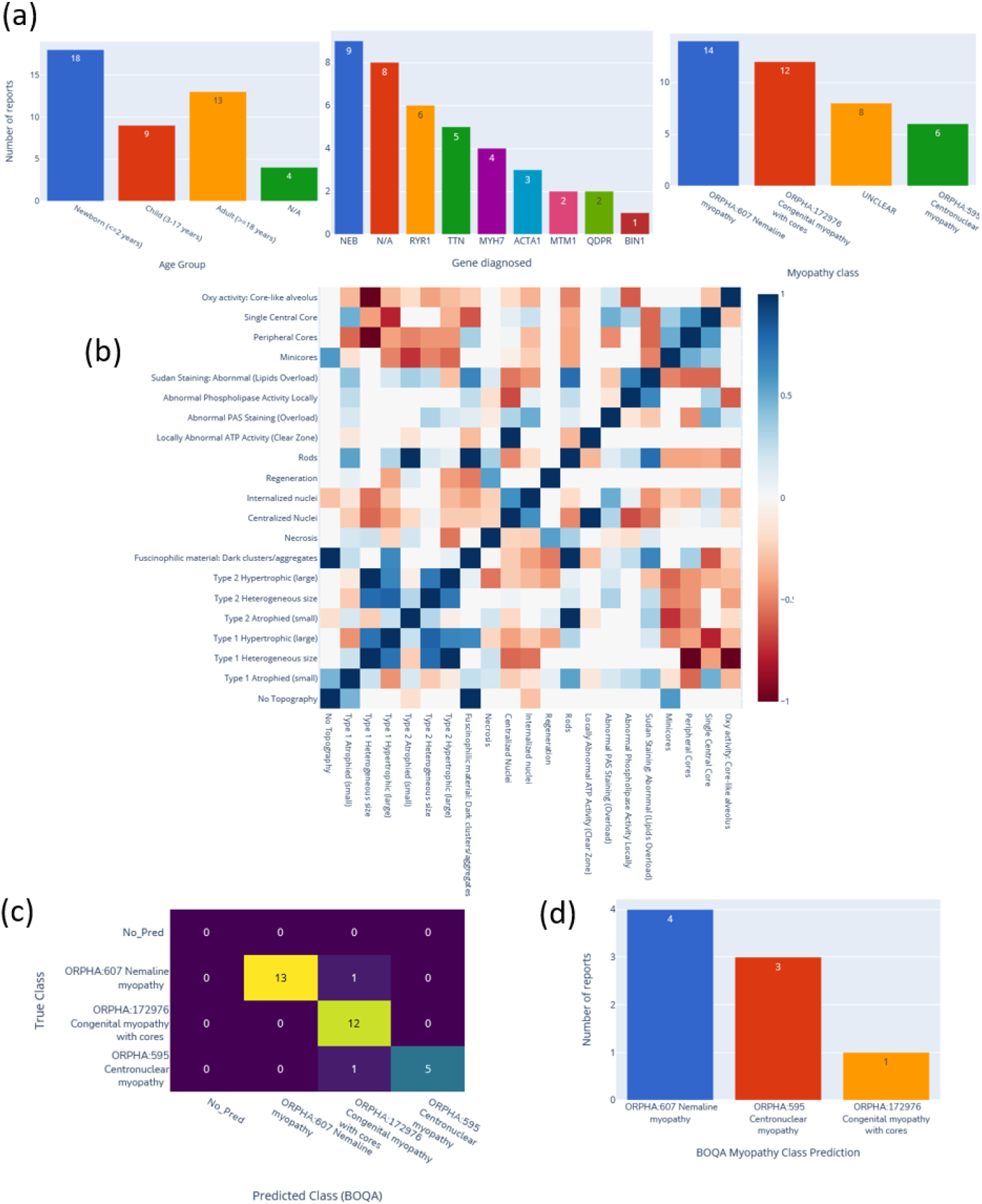
Automatic visualization of 40 generated congenital myopathy reports. **(a)** Histogram of the number of reports by age group, by diagnosed gene (top 9) or by congenital myopathy class. **(b)** Correlation matrix of standard vocabulary terms after annotation for all reports. **(c)** Confusion matrix of BOQA algorithm performance for suggestion of the three main congenital myopathy classes (NM, COM, CNM, n=32). Colors indicate the number of reports for each cell of the matrix, the lighter the color the more reports. **(d)** Histogram of the reclassification by BOQA of reports without a final diagnostic (n=8).

## DISCUSSION

IMPatienT is a platform that simplifies the digitization, processing, and exploration of both textual and image patient data. The web application is centered around the concept of a standard vocabulary tree that is easy to create and used to process text and image data. This allows IMPatienT to work with patient data from domains that still lack a consensus ontology and rely on well-established ontologies for patient data, such as HPO for phenotypes, Orphanet for disease names or HGCN/HGVS for genetic diagnoses.

The semi-automatic approach implemented in IMPatienT offers faster digitization processes while ensuring accuracy through manual review. This is achieved by analyzing text data using OCR and NLP to automatically match the text to the standard vocabulary, followed by manual correction. For image data, the user first provides sparse annotations on the image, which are then used to compute an automatic segmentation of the whole image. For data exploration, IMPatienT uses a fully automatic approach including various visualizations as well as diagnosis suggestions, while allowing the user to extract the processed data in a standard format for further analysis (database, images, frequency tables).

IMPatienT aims to integrate multiple approaches in a unified platform with two main objectives: universality (*i*.*e* not restricted to a specific domain) and multimodality (*i*.*e*. integration of multiple data types). To our knowledge, other tools similar to IMPatienT do not fulfill both objectives.

We performed a comparison of the main functionalities of IMPatienT with other tools used in the community. Phenotips, SAMS and PhenoStore are similar to IMPatienT as they are designed as a patient information database. However, they are restricted to processing patient phenotype data by using HPO and do not integrate multimodal data. IMPatienT goes further by allowing for custom observations with the vocabulary builder and with automatic digitization with OCR/NLP as well as integrating tools to exploit image data.

Other tools are similar to one or two modules only of IMPatienT. For example, Doc2HPO is a tool that also uses a semi-automatic approach to digitize clinical text according to a list of HPO terms, based on NLP methods and negation detection. However, as Doc2HPO is also restricted to HPO, it does not provide custom vocabulary tree facilities. In contrast IMPatienT is suitable for digitization of text data from any domain of interest.

For image data, software such as Cytomine and Ilastik are widely used and perform well on biological data, but they do not allow the user to take into consideration the multimodal aspects of patient data by keeping the raw image and the expert interpretation (histological report) in a single database along with a collaborative and rich-defined custom ontology.

Finally, in IMPatienT we reimplemented the diagnosis suggestion algorithm called BOQA that is also used in Phenomizer, a tool to rank a list of the top matching diseases based on a list of input HPO terms. We modified the algorithm to consider frequencies of terms by disease to have meaningful predictions. However, BOQA uses binary states for terms (terms are marked as present or absent) and is not compatible with numeric features. In the future, it will be necessary to implement a more complex system such as explainable AI with learning classifier systems (Urbanowicz & Moore, 2015). This should improve accuracy, explainability, and handling of quantitative values, although at the cost of computational power.

IMPatienT still lacks some feature compared to other tools, such as a pedigree editor, support for DICOM and gigapixel images and phenotypic data export to the Phenopacket format. In the future, we plan to further develop IMPatienT by adding these features to the interface. We also want to explore the automatization of the standard vocabulary creation with the analysis of a complete corpus of text. For text analysis, we wish to implement additional context comprehension, *i*.*e*. not only negation but also hypothetical statements, uncertainty and family context as well as better text-vocabulary terms matching. Finally, we plan to expand the scope of the OCR/NLP method by integrating existing NLP tools to automatically detect HPO terms, gene symbols and disease name the report text.

With IMPatienT, we have developed an integrated web application to digitize, process and explore multimodal patient data. IMPatienT can serve as a research tool to find new associations of patient features that might be relevant for diagnosis. A demonstration instance of the web application is available at https://impatient.lbgi.fr.

## Source Code and Data Availability Statement

The source code for IMPatienT v1.5.0 is available in its GitHub repository (https://github.com/lambda-science/IMPatienT). The synthetic dataset generated and analyzed during the current study is also available in the same repository.

## Acknowledgements

We thank the BiGEst-ICube platform for their assistance. We thank the Agence Nationale de la Recherche (ANR), 80 | Prime CNRS (MYO-xIA Project), the University of Strasbourg and INSERM for funding this work.

## Supplementary Materials

- **Table S1** - table_frequencies_per_gene.pdf - **Table of frequencies of standard vocabulary per genes**. This PDF file contains all frequencies of standard vocabulary terms for each gene with the total number of reports per gene and the number of occurrences of each term if not 0.
- **Table S2** - table_frequencies_per_diag.pdf - **Table of frequencies of standard vocabulary per diagnosis**. This PDF file contains all frequencies of standard vocabulary terms for each diagnosis with the total number of reports per diagnosis and the number of occurrences of each term if not 0.

**Supplementary Figure S1:**
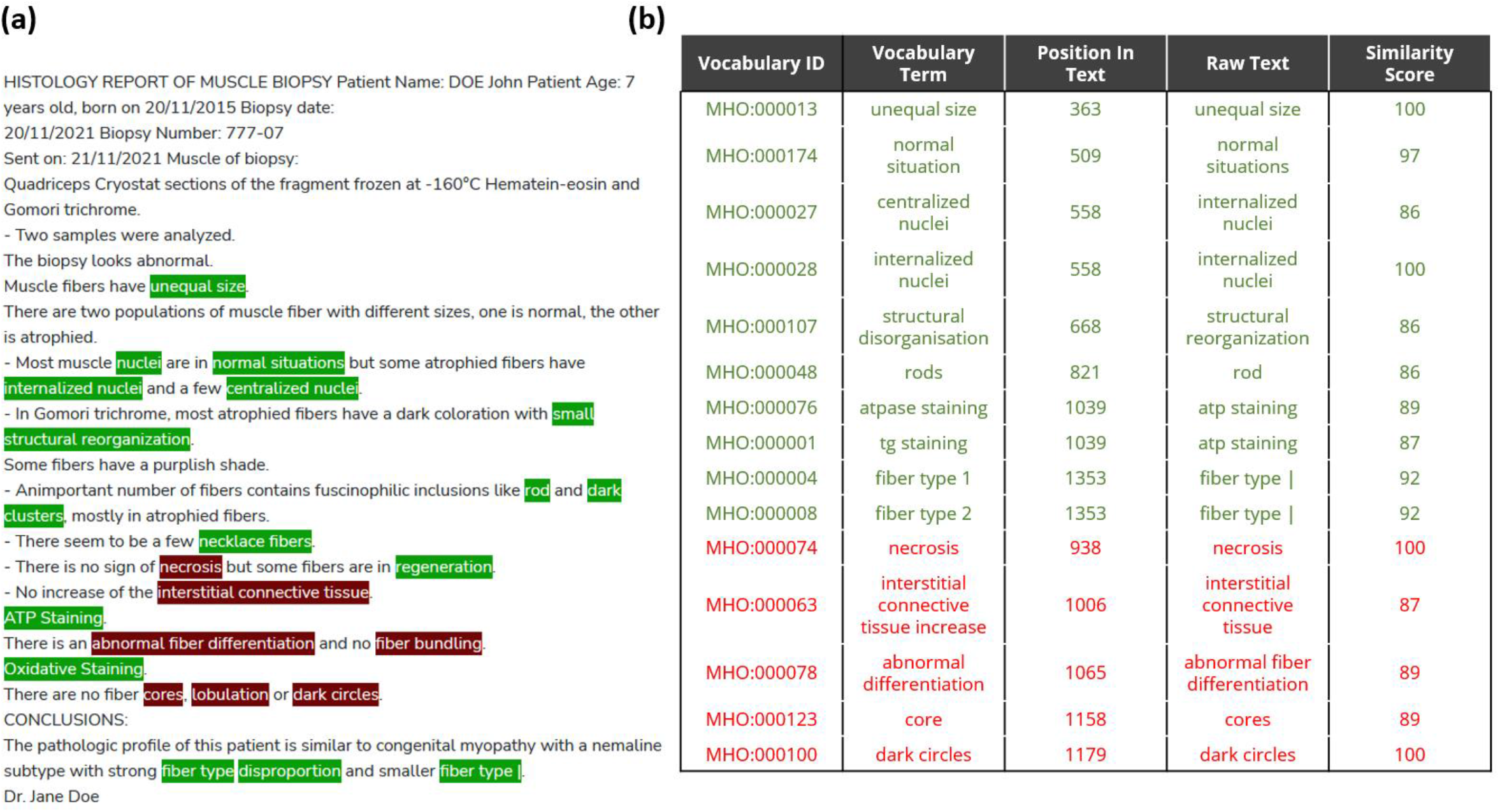
Qualitative assessment of the performances of the NLP method matching text to the standard vocabulary. (**a)** Raw muscle histology report text with detected keywords highlighted in green and red. A red highlight indicated that the keyword is in a negated sentence. **(b)** Table of some highlighted keywords and the details of the match (matching vocabulary ID and terms, position in the raw text, matching n-gram [raw text] and the similarity score of the comparison). Green and red colors correspond to keywords detected as present and present in negated sentence respectively.

